# Light Sheet Theta Microscopy for High-resolution Quantitative Imaging of Large Biological Systems

**DOI:** 10.1101/119289

**Authors:** Bianca Migliori, Malika S. Datta, Christophe Dupre, Mehmet C. Apak, Shoh Asano, Ruixuan Gao, Edward S. Boyden, Ola Hermanson, Rafael Yuste, Raju Tomer

**Affiliations:** Department of Biological Sciences; NeuroTechnology Center; Data Science Institute Columbia University; Department of Neuroscience, Karolinska Institut, Sweden; MIT Media Lab and McGovern Institute, Departments of Biological Engineering and Brain and Cognitive Sciences, MIT, Cambridge, MA USA

## Abstract

Advances in tissue clearing and molecular labelling methods are enabling unprecedented optical access to large intact biological systems. These advances fuel the need for high-speed microscopy approaches to image large samples quantitatively and at high resolution. While Light Sheet Microscopy (LSM), with its high planar imaging speed and low photo-bleaching, can be effective, scaling up to larger imaging volumes has been hindered by the use of orthogonal light-sheet illumination. To address this fundamental limitation, we have developed Light Sheet Theta Microscopy (LSTM), which uniformly illuminates samples from same side as the detection objective, thereby eliminating limits on lateral dimensions without sacrificing imaging resolution, depth and speed. We present detailed characterization of LSTM, and show that this approach achieves rapid high-resolution imaging of large intact samples with superior uniform high-resolution than LSM. LSTM is a significant step in high-resolution quantitative mapping of structure and function of large intact biological systems.

## Introduction

The emergence of various tissue clearing and molecular labelling methods over the last decade is enabling unprecedented optical access to the structure and function of intact biological systems (Becker et al., 2012; Chung et al., 2013; Dodt et al., 2007; Erturk et al., 2012; Hama et al., 2011; Ke et al., 2013; Kuwajima et al., 2013; Murray et al., 2015; Pan et al., 2016; Renier et al., 2014; Romanov et al., 2017; Susaki et al., 2014; Tomer et al., 2014; Yang et al., 2014). Most of these methods employ a cocktail of chemicals for membrane lipid dissolution and/or refractive index smoothening to render the tissue transparent (Richardson and Lichtman, 2015). Together with parallel advances in high-speed microscopy methods, these approaches have already proven to be highly effective in mapping of organs as large as the intact adult mouse brain (Lerner et al., 2015; Migliori et al., 2016; Romanov et al., 2017; Tomer et al., 2014). By providing a highly-detailed 3D view of the architecture of normal and abnormal intact tissues, these methods can accelerate our understanding of the structure and function of brains, a key goal of high profile BRAIN initiative, as well as provide mechanistic insights into the pathophysiology of the microarchitecture of diseased tissues. However, scaling up these approaches, while maintaining uniform high imaging quality, faces the challenges of clearing and labelling of large samples combined with high-resolution quantitative 3D imaging.

Here we address some of these challenges by developing a conceptually distinct microscopy framework: Light Sheet Theta Microscopy (LSTM). Building upon the principles of Light sheet microscopy (LSM)(Siedentopf and Zsigmondy, 1903; Stelzer, 2015), LSTM allows high-speed quantitative imaging of large intact tissues at uniform high resolution. The basic concept of LSM was first introduced more than a century ago (Siedentopf and Zsigmondy, 1903). LSM uses a thin sheet of light for planar illumination of a sample and an orthogonally arranged wide-field detection arm for simultaneously capturing the emitted signal with a high speed CCD or sCMOS camera (Migliori et al., 2016). Compared to other commonly used 3D imaging modalities, confocal and 2-photon microscopy, LSM places the minimum possible energy load on the sample and provides orders of magnitude faster imaging. LSM has been highly successful for experimentations in developmental biology (Huisken et al., 2004; Keller et al., 2008; Preibisch et al., 2010; Reynaud et al., 2015; Wu et al., 2013), cell biology (Chen et al., 2014; Gao et al., 2012; Planchon et al., 2011), high-resolution whole brain neuroanatomy (Lerner et al., 2015; Tomer et al., 2015; Tomer et al., 2014) and neural activity mapping experiments (Ahrens et al., 2013; Bouchard et al., 2015; Chhetri et al., 2015; Holekamp et al., 2008; Tomer et al., 2015; Vladimirov et al., 2014). The samples larger than the field-of-view (FOV) of a microscope are imaged by sequential acquisition of overlapping image stacks (Tomer et al., 2014), which are then computationally stitched to result in the final image volumes.

The sizes of samples that can be imaged with LSM is restricted along two dimensions, the detection and the illumination axes (**Figure 1**). Sample illumination is therefore restricted to few millimeters deep and wide, without a limit on length. Although LSM has been used for rapid high-resolution imaging of samples as large as mouse brains (Renier et al., 2014; Susaki et al., 2014; Tomer et al., 2014), image quality progressively reduces towards the center because of illumination light scattering (even with two-sided illumination). The reduction of image quality is further more severe for larger rat brain tissues(Stefaniuk et al., 2016). These fundamental limitations have precluded the use of LSM for high-resolution quantitative imaging of large samples such as rodent brain tissues, and for imaging of a sample with laterally extended geometries such as thick slices of human brain or physically expanded tissues (e.g. using Expansion Microscopy(Chen et al., 2015)). Recently, alternative optical configurations of LSM have been developed that partially address these limitations, including the rotation of the illumination and the detection axes by 45° relative to the sample surface normal as done in OCPI, iSPIM and diSPIM implementations (Holekamp et al., 2008; Strnad et al., 2016; Wu et al., 2011; Wu et al., 2013) and the generation of illumination light sheets through the detection objectives itself as implemented in SCAPE/OPM (Bouchard et al., 2015; Dunsby, 2008). iSPIM/diSPIM approach removes limits on the lateral dimensions of imaging volumes, although at the cost of reduced effective working distance of the detection objective, and therefore is restricted to relatively low-resolution detection for thick samples (**Figure 1a**). SCAPE/OPM approaches employ rotation optics to image an oblique plane in a sample, illuminated through the detection objective itself. This configuration enables fast volumetric imaging speeds for small volumes, although at the cost of reduction in image quality due to the collection of signal from non-native focal planes (**Figure 1a**). While these implementations have been highly successful for rapid imaging of small samples (e.g. *C. elegans* embryos) or small volumes of mouse brain cortex, they suffer from other geometric and image quality constraints when applied to larger samples.

**Figure 1.**
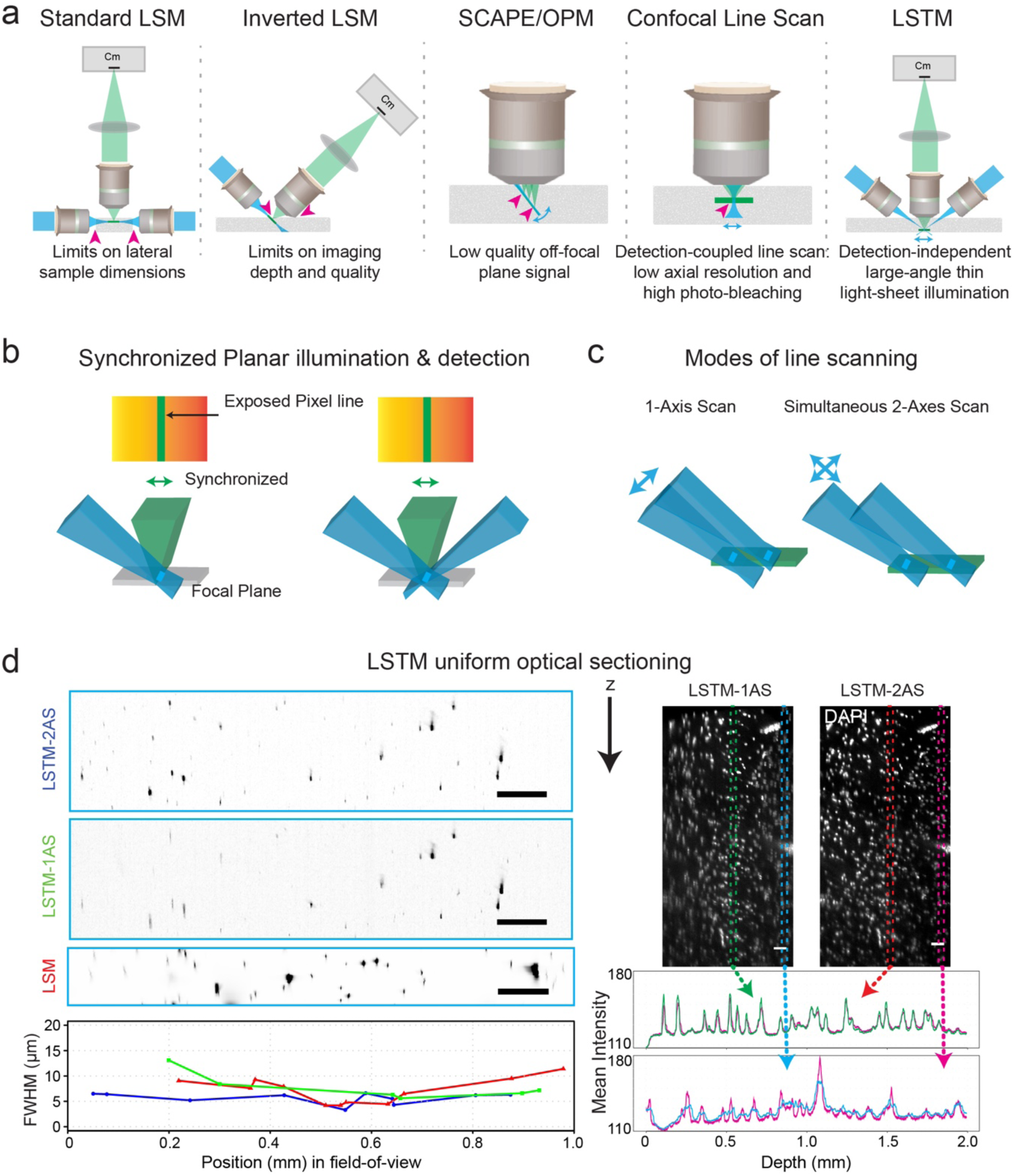
Light Sheet Theta Microscopy (LSTM) for high-resolution quantitative imaging of large intact samples. (a) Schematic comparison of Light Sheet Microscopy (LSM), inverted LSM, SCAPE/OPM, Line Scan Confocal and Light Sheet Theta Microscopy (LSTM). LSM involves the illumination and the detection optics to be orthogonally arranged, thus putting limitations on the sample size along two of the sample dimensions i.e. along the detection and the illumination axis. Alternative implementations such as iSPIM, SCAPE/OPM and Line Scan Confocal are partially effective in addressing this for smaller samples, however at the cost of reduction in the effective detection working distance (magenta arrowheads) and image quality (for example, SCAPE collects low quality signal from non-native focal planes, and Line Scan Confocal results in poor axial resolution and high photo-bleaching) needed for quantitative imaging of larger samples. The LSTM configuration, which involves illumination through non-orthogonal axes (<90°), can effectively address these limitations, while providing high imaging speed and uniform resolution across large field-of-view, although at some cost of increased photo-bleaching for small samples (**Figure 3**). (b) Schematic representation of line detection strategy for optical sectioning. One or two thin sheets of light intersect with the detection plane in a line, which is then synchronously scanned (along with the rolling shutter of sCMOS camera) in the focal plane, resulting in uniform planar imaging. (c) Schematic summary of two modes of LSTM line scanning. The 1-Axis scanning (1-AS) involves the translation of the light sheet perpendicular to their propagation direction, whereas the simultaneous 2-Axes scan (2-AS) involves the translation perpendicularly as well as along the illumination direction such that the thinnest part of the sheet intersects with the detection plane. The latter mode results in uniform axial resolution across the entire field-of-view. (d) Comparison of optical sectioning in LSTM 1-AS, LSTM 2-AS and LSM configurations. Left shows x-z projection of 1 micron fluorescent beads imaged with LSTM 1-AS, LSTM 2-AS and LSM using the same detection objective (Olympus 10X/0.6NA/8mmWD). The illumination and detection objectives in LSTM configuration were arranged at ∼60° angular separation (as discussed in **Figure 3**). Axial FWHMs (Full Width at Half Maximum) were calculated from beads positioned at different positions along the field-of-view and plotted in the bottom graph with corresponding colors (blue: LSTM in 2-AS mode, Green: LSTM in 1-AS mode, red: LSM). LSTM 2-AS achieves uniform optical section over the entire field-of-view whereas both LSTM in 1-AS mode and LSM results in similarly reduced axial resolution on the peripheries. Right shows the x-z projections (20 microns thick) of an image stack acquired from a DAPI stained CLARITY-cleared Human brain slice, using LSTM 1-Axis and LSTM 2-Axes scan modes with 10X/0.6NA/8mmWD (Olympus) detection objective. The graphs below compare the signal along the imaging depth for a central and a peripheral region-of-interest. This example demonstrates that LSTM enables the use of very thin light sheets for achieving good axial resolution over a large field-of-view. All scale bars are 100 microns. See also **Supplementary Figure 1** for a comparison of different camera exposure settings, and **Supplementary Video S1** for 3D reconstructions of the complete stacks.

We have developed Light Sheet Theta Microscopy (LSTM) to address some of these limitations. LSTM achieves planar imaging by employing obliquely arranged illumination light sheets from the same side of the sample as the detection objective. This configuration alleviates limitations on the lateral dimensions of the sample, while providing similar imaging depth, uniform high-resolution and low photo-bleaching (higher than LSM for smaller samples, but similar to LSM for larger samples). Note that LSTM bear some resemblance to Line Scan Confocal Microscopy (LSCM) (Mei et al., 2012; Wolleschensky et al., 2006) at an abstract level, as effectively an illumination line profile is scanned in the detection focal plane (**Figure 1a**). In LSCM, the illumination line profile is generated by focusing of a collimated LASER beam through the detection objective itself, which is then rapidly scanned over the focal plane to result in high imaging speed(Wolleschensky et al., 2006). However, this increase in imaging speed comes at the cost of reduction in imaging quality (both axial and lateral resolution) and very high photo-bleaching (similar to Confocal). Moreover, the usage of a single objective for illumination as well as for signal detection reduces the flexibility needed for using optimal optics for efficient illumination and high resolution imaging (which has been a critical factor in the success of LSM). These and other limitations make LSCM less suitable for high-resolution imaging of large samples. Indeed, LSCM has only be successfully used for relatively low resolution imaging of very small samples(Mei et al., 2012; Wolleschensky et al., 2006). The key advantages of LSTM approach over LSCM includes high imaging resolution and low photo-bleaching, both due to the use of thin oblique illumination light sheets, generated using two separate illumination objectives, making LSTM ideally suited for rapid high resolution imaging of very large samples. We present detailed characterization of the LSTM approach using examples that include mouse and rat brain, as well as human brain slices. Furthermore, we demonstrate the unique suitability of LSTM for rapid volumetric imaging of highly motile animals. Through high-speed quantitative imaging of larger samples, LSTM could facilitate mapping of an entire post-mortem human brain (slab-by-slab) in a practical time-frame.

## Results

### Light Sheet Theta Microscopy (LSTM)

LSTM includes a standard wide-field detection arm and two symmetrically arranged non-orthogonal (θ<90°, relative to the sample surface normal) illumination arms for the generation of thin sheets of light that intersect at the detection focal plane (**Figure 1,2**). This approach results in a thin line illumination profile which is then scanned, in synchrony with the line-by-line rolling shutter detection of an sCMOS camera (virtual slit effect, Tomer et al 2014), to achieve thin optical sectioning (**Figure 1**). In contrast to LSM, the non-orthogonal optical configuration of LSTM does not place any restrictions on the lateral dimensions of the imaging volume, while still allowing access to the complete working distance of the detection objective, provides high imaging speeds (20 milliseconds per image acquisition, same as COLM implementation(Tomer et al., 2014)) and similarly low photo-bleaching for large samples (**Figure 3d-f, Supplementary Figure 5**). To achieve planar illumination, we designed and tested two modes of line scanning: (1) 1-axis scanning (1-AS), which involves translation of light sheets perpendicular to their propagation direction (**Figure 1c-left)**. (2) simultaneous 2-axes scanning (2-AS): concurrent translation of light sheet perpendicular to and along the propagation direction so that the thinnest part of the light sheet intersects with the detection plane (**Figure 1c-right**) (note that similar and other approaches have been utilized for uniform illumination of a single field-of-view in LSM (Dean et al., 2015; Fu et al., 2016; Gao, 2015; Planchon et al., 2011; Tomer et al., 2015; Vettenburg et al., 2014)). The LSTM 1-AS approach provides a simpler implementation, although at the cost of non-uniformity in the planar illumination and low axial resolution (because of the need to use relatively low numerical aperture objectives for illumination). The 2-AS approach allows for uniform planar illumination and detection to enable high-resolution quantitative imaging. To characterize the two LSTM modes and to compare them with LSM, we imaged micron-sized fluorescent beads and CLARITY-cleared (Tomer et al 2014) human brain tissue stained with nuclear marker DAPI (**Figure 1d, Supplementary Figure 1, Supplementary Video 1**). The resulting image volumes reveals that LSTM 2-AS indeed allows for uniform high axial resolution across the entire field-of-view, whereas both LSTM 1-AS and LSM imaging produce reduced image quality on the periphery of the field-of-view (**Figure 1d, Supplementary Figure 1 and Supplementary Video 1)**. Simultaneous two-sided illumination (**Figure 1b-right**), from two symmetrically arranged illumination arms, provides higher signal and may reduce the illumination artifacts caused by opaque objects in the illumination path thus improving the uniform planar illumination and detection pre-requisite for achieving quantitatively accurate imaging of large transparent samples.

**Figure 2.**
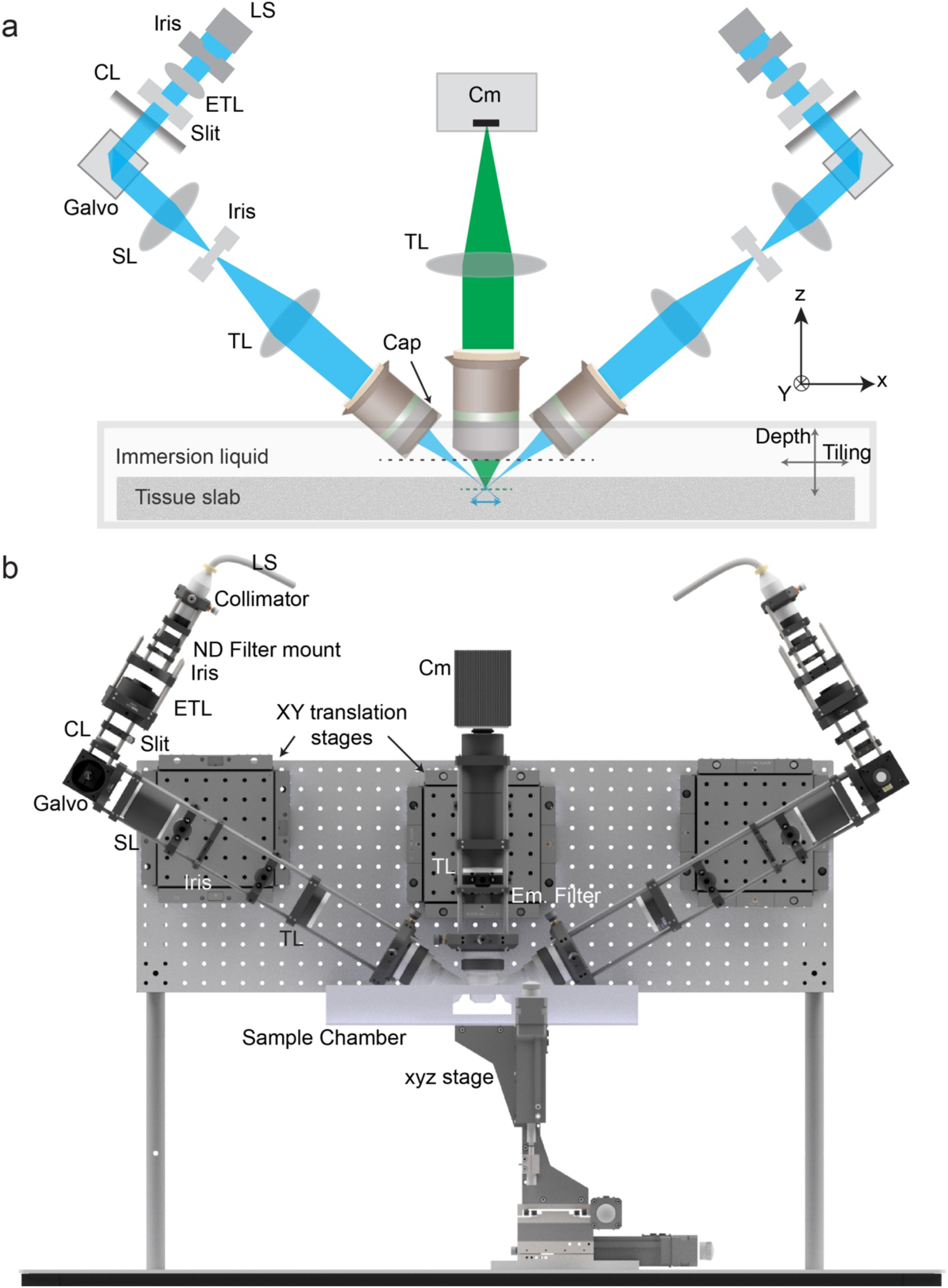
LSTM microscopy implementation. (a) Schematic summary of the optical path of LSTM microscopy. Two light sheets are generated by using a cylindrical lens (CL), scan lens (SL), tube lens (TL) and illumination objectives (Olympus Macro 4x). The galvo scanner is used to translate the light sheets perpendicular to their propagation direction, and the electrically tunable lens (ETL) is used to translate the thinnest part of the sheet along the propagation direction. Custom made collimators were used to generate an input beam of ∼10 mm diameter, which is then trimmed through an iris. A slit is placed after the ETL to reduce the beam diameter in one direction in order to control the effective numerical aperture of the illumination. An additional Iris is placed at the conjugate plane between the SL and TL to control the light sheet height for minimizing any unwanted illumination. The illumination axes are arranged at an approximate angle of 60° with the detection axis. A custom 3D-printed cap with a quartz coverslip was attached to the illumination objective (designed for imaging in air). The detection arm consists of the detection objective (Olympus 10x/0.6NA/8mmWD or 25x/1.0NA/8mmWD), the tube lens and a sCMOS camera (Hamamatsu Orca Flash V3.0). (b) 3D model of the entire LSTM microscope. A vertical breadboard was used to mount the optical components, via x-y translation stages to allow for degree of freedom during the optical alignment. A sample chamber was mounted on a 3 axes (x,y,z) motorized stage. Samples were mounted in a quartz cuvette of appropriate size, which was then attached at the bottom of the sample chamber. Imaging of the entire sample was performed by moving the sample chamber. See also **Supplementary Figure 2** for further details of the microscope, **SupplementaryVideo 2** for an animation of the 3D model and **Supplementary Table 1** for complete parts list.

### LSTM implementation

A standard LSM system consists of a fixed orthogonal arrangement of illumination and detection optical arms that are typically positioned using independently mounted optical components. For example, CLARITY-Optimized Light-sheet Microscopy (COLM, Tomer et al 2014) uses a horizontally spread out configuration in which each component is free to move independently. The need to maintain degrees of freedom on translation as well as rotation of the entire illumination assemblies as a whole poses challenges, particularly during fine adjustments of optical alignment. To implement the illumination arms as a rigid monolithic assembly that can be easily rotated and translated, we built the first prototype on a vertically mounted breadboard (**Figure 2, Supplementary Figure 2**), using a caging system to connect all the optics to rigid frames. The entire assemblies were then connected to the breadboard via xy manual translation stages to allow for finer positioning adjustments. We also designed an open top sample mounting strategy by employing a 3D printed chamber (**Supplementary Figure 2**) attached to a high accuracy x-y-z motorized stage assembly. Biological samples were mounted in a quartz cuvette of the appropriate size, tightly connected to the bottom of the sample chamber (**Supplementary Figure 2**). We also developed adapters to mount a prism mirror for the optical alignments. The entire sample chamber assembly can be translated in 3 dimensions to acquire the image volumes. This approach allows for full exploration of various parameters of the system (such as the angular separation between the illumination and detection arms) and acquiring data from large samples by providing rigid monolithic illumination and detection units with translational and rotational degrees of freedom.

The final overall LSTM illumination configuration includes a LASER source, collimators (∼10 mm output beam diameter), ETL, cylindrical lens, galvo scanner, scan lens, tube lens and illumination objective (**Figure 2a**). In addition, we incorporated an iris, after the collimator, to remove the peripheral spread of Gaussian beams, a one dimensional slit, before cylindrical lens, to control the effective numerical aperture of illumination and a second iris at the conjugate plane, between scan lens and tube lens, to control the light sheet height. The detection arm is composed of a detection objective, emission filter, tube lens and an sCMOS camera.

Since LSTM involves scanning of a line illumination-detection profile generated by the intersection of the light sheet and the detection plane, we used static sheets (generated by the use of a cylindrical lens and associated optics), instead of a dynamic sheet (generated by rapid scanning of a pencil beam) to maximize the imaging speed. The cropping of peripheral parts of the large input diameter beam with an iris ensured a relatively uniform intensity distribution profile across the static light sheet. We used a galvo scanner to achieve rapid translation of light sheets perpendicular to their propagation direction. Finally, for the 2-AS mode, we also needed rapid translation of the thinnest part of the sheet along the propagation direction. Possible approaches here include fast piezo motors to translate the illumination objectives, using holographic spatial light modulators or an electrically tunable lens (ETL) able to drive the divergence and convergence of a collimated beam. The use of piezo motors for rapid scanning of objective often results in vibrations and require additional settling time(Ahrens et al., 2013; Tomer et al., 2015), and the spatial light modulators are limited in modulation speed because of slower refresh rates. ETLs, on the other hand, can achieve high frequency modulation of focal point position without the need for moving optics of significant mass(Fahrbach et al., 2013). We thus tested an ETL based approach and found it to be highly effective for achieving uniform simultaneous 2-axes scanning (**Figure 1d, Supplementary Videos 1**).

The LSTM assembly was optically aligned by placing a prism mirror (with fine scratches in the center, see **Supplementary Figure 2** for mounting arrangements) in the focal plane of detection optics, to visualize the location and cross-section of the light sheet relative to the detection focal plane. The light sheet positioning parameters were optimized such that the thinnest part was in alignment with the center of the field-of-view of the detection plane. Next, the mirror was replaced with a high concentration (>2%) agarose gel containing fluorescent beads (Note, the high concentration of agarose was used to ensure that only the surface plane of the agarose gel was visible during the alignment optimization.). Optimal galvo scanner and ETL parameters for achieving uniform planar illumination across the entire field-of-view were identified by examining the extent and quality of the illuminated beads located on the surface.

### LSTM characterization

A series of calculations were used to assess and compare various properties of LSTM (summarized in **Figure 3**). First, we devised a method to calculate the physical geometric constraints of arranging a given set of detection and illumination objectives in a non-orthogonal configuration (**Supplementary Figure 3**). The main physical parameters used in the calculations are the working distances and the diameters of both the illumination and detection objectives. We calculated the range of physically-allowable, relative angular arrangements that enable light sheets to intersect the detection focal plane at their thinnest parts, while also ensuring that illumination objectives remain above the physical extent of the detection objective (**Supplementary Figure 3**). For instance, only angular configuration of 43-62 degrees for Macro 4x/0.28NA/29.5mmWD (Olympus) as the illumination objective and 10x/0.6NA/8mmWD (Olympus) as the detection objective are possible (**Figure 3, Supplementary Figure 3)**. Note that the working distance of this illumination objective is given for use in air, and therefore we calculated the approximate effective working distance as shown in **Supplementary Figure 3**. Next, determined the influence of angular separation of illumination and detection arms on the resulting image volumes. We first calculated and compared the illumination path length in LSTM to LSM: shorter the illumination path length the better the image quality. In LSM, the illumination light sheet needs to penetrate the entire width of the sample for complete coverage, whereas in LSTM the effective illumination path length depends on the angular arrangement and the tissue thickness(t): *t/*cos(θ). As shown in **Figure 3b**, LSTM is expected to outperform LSM for high quality imaging of large samples. Next, we performed computer simulations to estimate the energy load (hence photo-bleaching) in LSTM vs. LSM imaging (**Figure 3d-f, Supplementary Figure 5**). The energy load redundancy in LSTM is expected to be independent of the sample’s lateral dimensions, whereas it increases with the sample width in LSM (as the light-sheet needs to travel through the entire width of the sample for every acquired image in a given tile, **Figure 3d**). **Figures 3e** summarizes the effects of various system parameters (sample width, angular separation of illumination and detection axes, sample thickness, and detection objective magnification) on the overall energy load in LSTM and LSM. It is evident that LSTM imparts significantly higher energy load, compared to LSM, for imaging of smaller samples, while similar for larger samples. The LSTM energy load also decreases when using higher angular configuration (between illumination and detection axes) and higher magnification objectives. Nevertheless, we provide strong empirical evidence of no consequences in terms of photo-bleaching (summarized in **Figure 3f**), by imaging of fixed samples of various sizes and shapes, and also by high-speed long-term live functional imaging of a freely moving animal.

**Figure 3.**
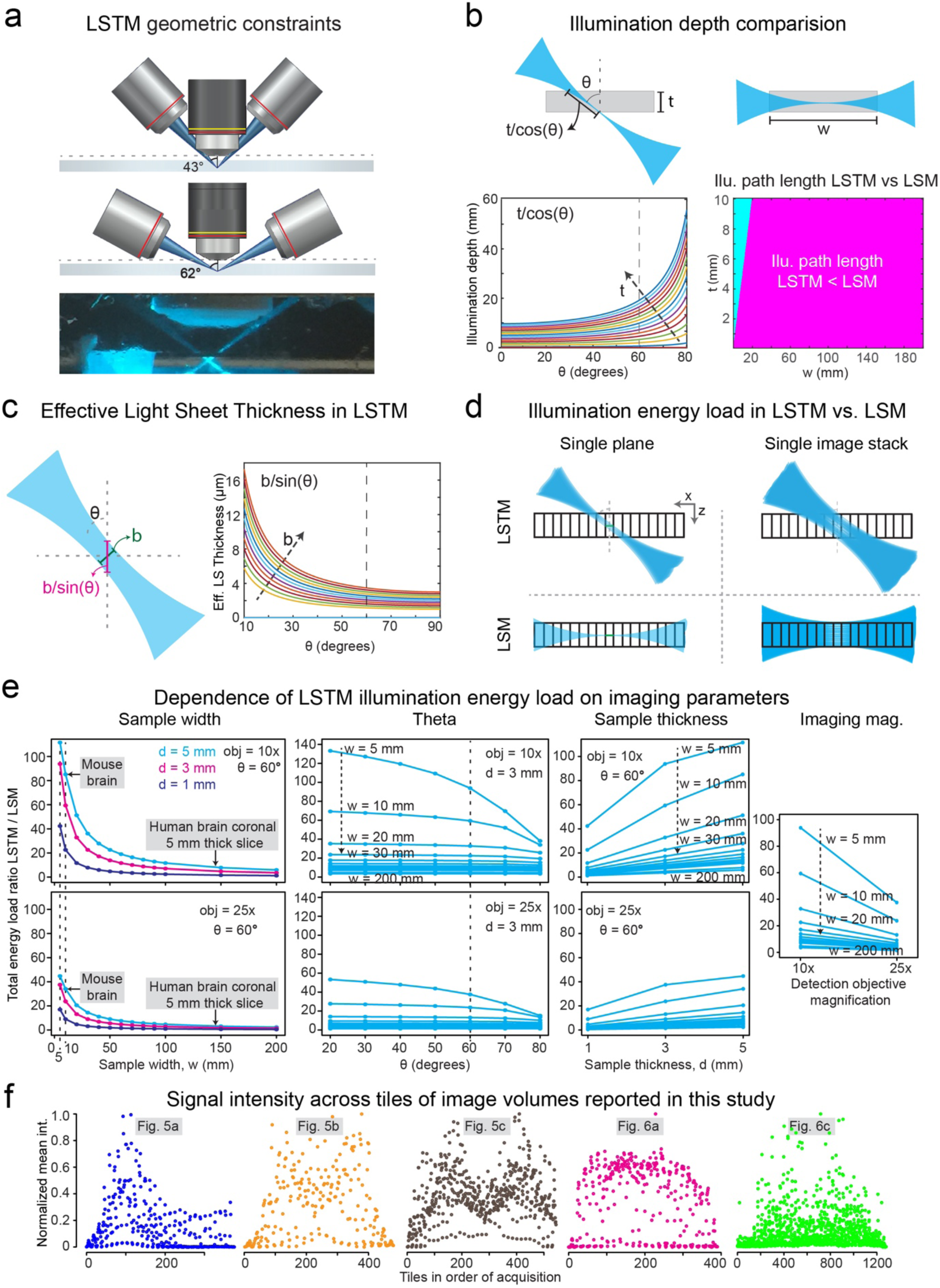
LSTM design parameters. (a) Physical geometric constraints for arranging the detection and illumination objectives. The schematics summarize the analyses for a particular set of objectives that were commonly used for LSTM imaging: Olympus Macro 4x/0.28NA/29.5WD for illumination, and Olympus 10x/0.6NA/8mmWD for detection. Two extreme angular positions are shown. Note that the specified working distance of the illumination objective (in air) was used to calculate the elongated working distance (owing to the high refractive immersion oil), as shown in **Supplementary Figure 3**. For all the imaging experiments, the angular separation between the illumination and detection optics was set to ∼60°. The bottom image shows an example of the physical implementation of generating illuminating thin sheets. (b) Comparison of the illumination path length in the LSTM and LSM. In LSM, the illumination light sheet will need to penetrate the entire width of the sample, whereas in LSTM it is dependent on the angular arrangement and tissue thickness (t): *t/*cos(θ). The left graph plots the dependence of illumination depth of LSTM on θ. Each color curve represents a different tissue thickness(t), as marked by the arrow in the direction of increase in thickness. The right graph compares the illumination depth required to image a sample of given width and thickness. The ratio of the LSTM and LSM illumination path was converted into a binary representation to summarize the parameters ranges where LSTM (magenta) and LSM (cyan) need smaller illumination depth, and hence will provide better image quality. See **Supplementary Figure 4** for other angular configurations. (c) Effective planar illumination thickness can be approximated as b/sin(θ), where b is the light sheet thickness. The right graph plots the effective light sheet thickness as a function of θ for different values of b (the arrow points in the direction of increasing b value). (d) Comparison of energy load on the sample while imaging with LSM and LSTM. The two rows compare the time-accumulated energy load in LSM and LSTM for imaging of a single plane and a single image stack tile. The energy load in LSTM is expected to depend on the sample thickness for a given angular arrangement, and in LSM on the sample width. (e) Ratios of total illumination energy load is calculated as a function of systems parameters: sample width (w), angular configuration of LSTM (θ), sample thickness (d) and two detection objectives (10x/0.6NA/8mmWD and 25x/1.0NA/8mmWD). Illumination energy load is higher in LSTM for smaller samples and similar to LSM for wider samples. LSTM energy load decreases with angular separation between illumination and detection arms (60° is marked), and also with the magnification of detection objective. **Supplementary Figure 5** provides details of energy load calculations. (f) Average signal of tiles across entire image volumes is plotted for the datasets reported in this study. Note that no significant photo-bleaching is observed. See also **Supplementary Figures 3-5** for further detailed characterizations and calculation procedures.

In LSTM, from an illumination path length stand point, minimizing the angular separation will increase the imaging quality. However, when the effect of θ on the effective light sheet thickness (approximated as b/sin(θ), **Figure 3c**), which determines the axial resolution, is measured, an inverse relationship is found: the more the θ the better the axial resolution. Because illumination is provided via a relatively low NA objective (0.28) for which the light scattering has much smaller effect on the illumination side, we decided to maximize the angular separation (∼60°) to achieve higher axial resolution. All the experiments were performed using this configuration.

### LSTM enables rapid quantitative imaging of large samples with uniform high-resolution

We first tested the use of lower NA illumination (hence larger field-of-view and thicker sheet) in LSTM 1-AS configuration in a large sample, a CLARITY-cleared thick coronal section of *Thy1-eYFP* transgenic mouse brain (**Figure 4**). While, the LSTM 1-AS mode allowed for high-quality imaging of the section, image quality was reduced (marked with dotted-rectangles) for peripheral most portions of the field-of-view, even for the low NA illumination configuration. This result was similar to the imaging performance of a LSM system employing Gaussian beams for illumination.

**Figure 4.**
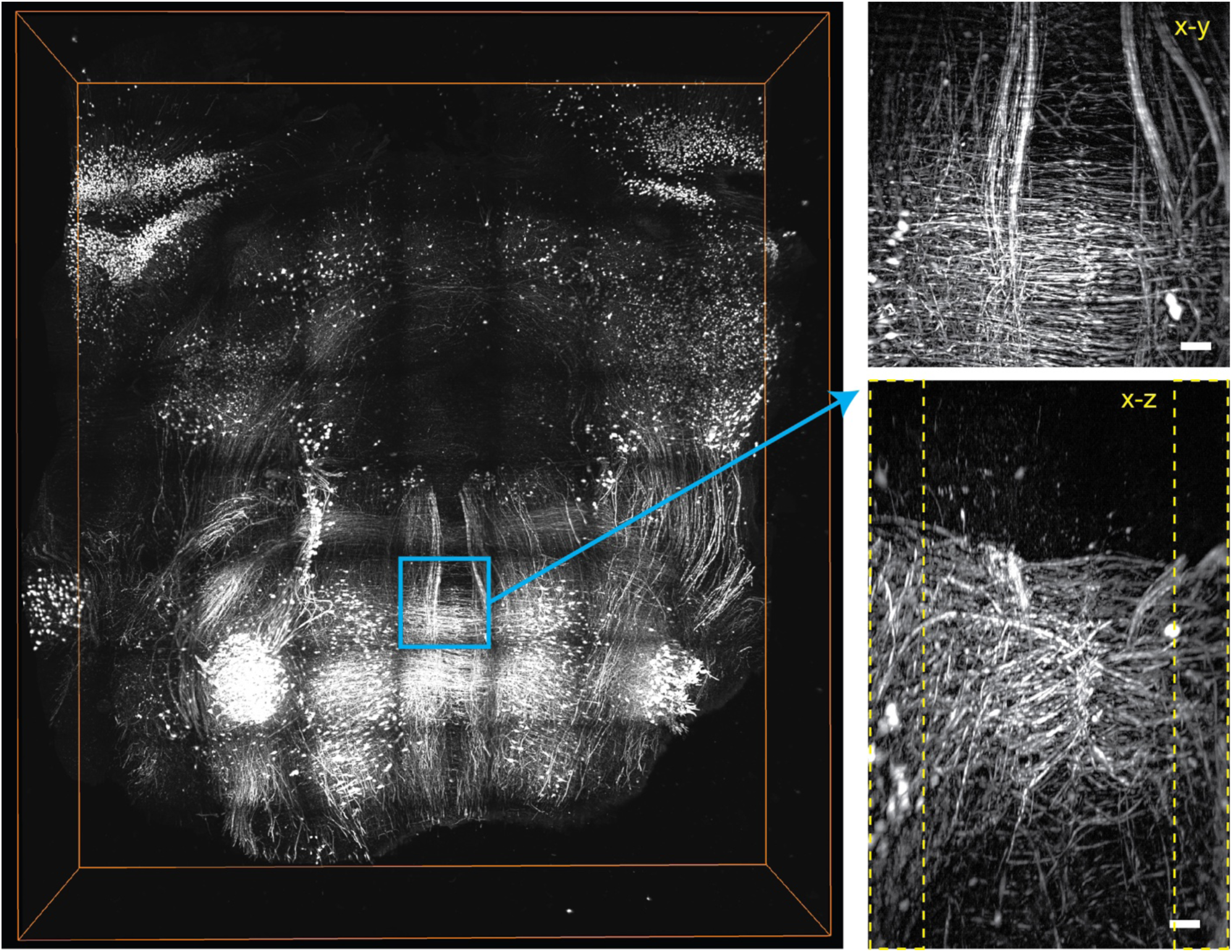
LSTM imaging quality in 1-AS mode. A thick slice of CLARITY-cleared *Thy1-eYFP* transgenic mouse brain was imaged with LSTM using 1-Axis scanning. Low numerical aperture illumination was used to generate a large field-of-view (and hence thicker) light sheets. The bounding box is 8.3mm x 7.3 mm x 2 mm. X-Y and X-Z maximum intensity projections of a stack at marked positions are shown on the right side. Similar to standard LSM, the image quality partially degrades on the periphery of the stack (outlined by the yellow dotted rectangles). The scale bar is 100 microns.

By allowing use of high NA objectives for illumination (hence thinner sheet), LSTM 2-AS mode enables high-resolution imaging with uniform quality. To assess the quantitative imaging performance of LSTM, we performed imaging of cleared intact mouse central nervous system of *Thy1-eYFP* transgenic mouse. As demonstrated in **Figure 5a** and **SupplementaryVideo 3**, LSTM enables rapid high-resolution quantitative imaging of these large samples without any reduction in the image quality across the sample dimensions. We further imaged a large (∼9.6mm x 13.5 mm x 5.34 mm) coronal slice of CLARITY-cleared *Thy1-eYFP* transgenic mouse brain, with 10x/0.6NA/8mmWD (**Figure 5b, Supplementary Video 4**) and 25x/1.0NA/8mm (**Figure 5c)** objectives, and larger input beam diameter (to employ the full available NA of 0.28) of the illumination objective. Note that this sample was expanded ∼1.5-2 fold by incubation in glycerol solution(Tomer et al., 2015) to result in ∼1-5-2 folds expansion. As demonstrated by zoom-in views of various locations of the samples, LSTM provides high uniform quality across the samples.

**Figure 5.**
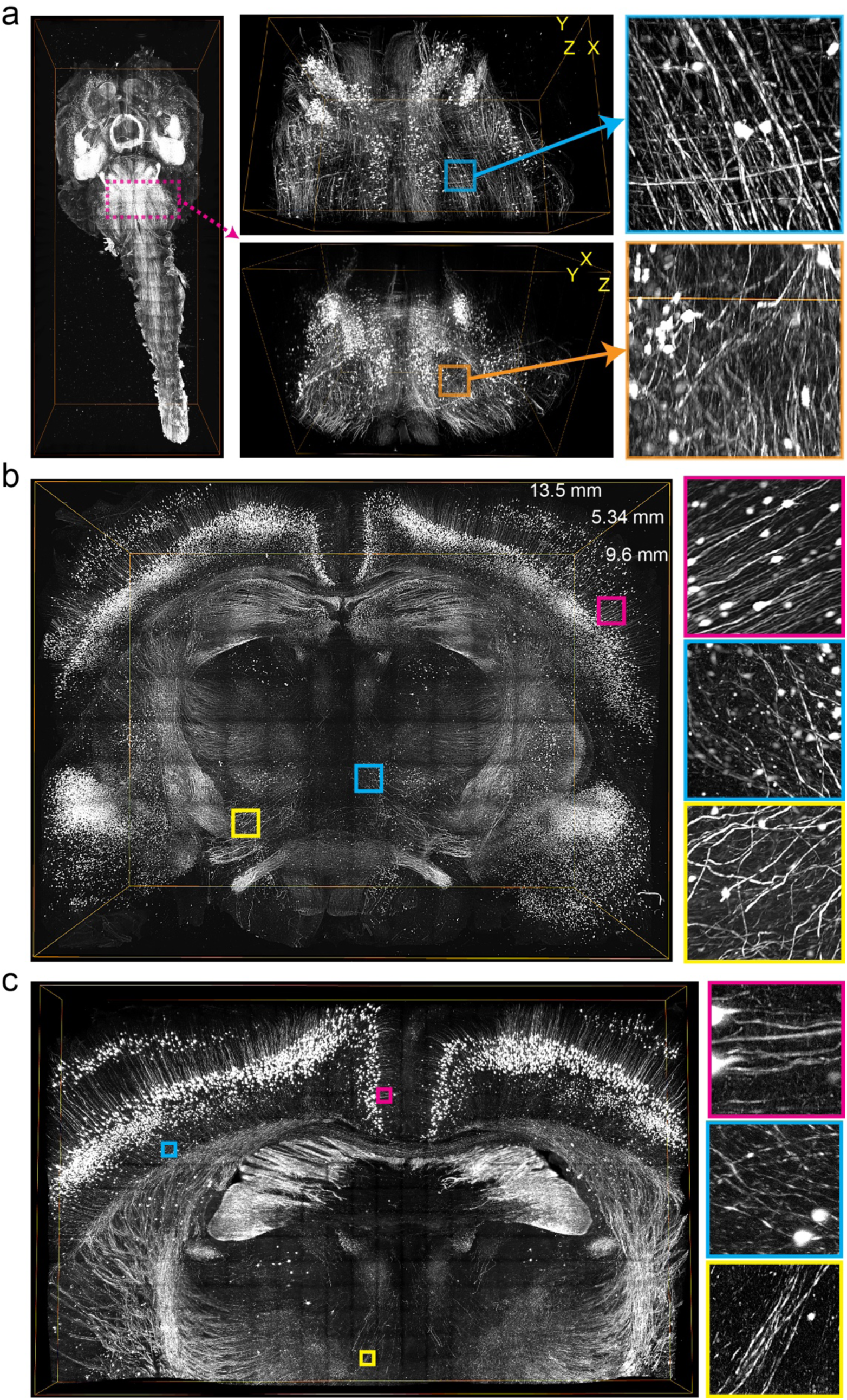
Rapid uniform high-resolution imaging of intact mouse central nervous system. (a) A CLARITY-cleared intact *Thy1-eYFP* transgenic mouse brain, with attached spinal cord, was imaged with LSTM microscopy (10x/0.6NA objective). High-resolution 3D rendering, using 2x2 fold down-sampled data, is shown for the entire tissue and for a sub-volume marked by magenta rectangles. Two orthogonal views, and zoomed-in images are shown as marked by corresponding colors. The bounding boxes are 11.8mm x 27.6 mm x 5.2 mm for the whole sample, and 5.1mm x 3.1mm x 3.5mm for the sub-volume. A detailed volume rendering is shown in **Supplementary Video 3**. (b) A large coronal slice of a *Thy1-eYFP* transgenic mouse brain was imaged with LSTM to demonstrate the uniform high quality imaging of LSTM. The volume rendering was performed using 4x4 fold down-sampled data. Zoomed-in images are shown for the marked colored squares, demonstrating image quality across different locations of the image volume. The bounding box is 9.6mm x 13.5 mm x 5.34 mm. **Supplementary Video 4** show detailed volumetric rendering. (c) The same sample was imaged with a high-NA 25x/1.0NA objective. The volume rendering was performed from 2x2 fold down-sampled images, and the zoomed-in images are shown at different locations across the sample, as marked by the colored squares. The bounding box is 6mm x 9.6 mm x 0.5 mm. See also **Supplementary Videos 3-4** for detailed volume renderings.

Next, we demonstrate that LSTM indeed outperforms LSM for uniform high-resolution imaging of large samples (**Figure 6**). For this we cleared a large slice of rat brain and stained it with a relatively uniform label to visualize all the blood vessels. Previous attempts of using LSM to image rat brain resulted in poor image quality apart from the periphery of the tissue(Stefaniuk et al., 2016), as also expected because of illumination scattering. We challenged the LSTM approach with this very large sample (∼2 centimeters wide and >5 mm deep) for a direct comparison with LSM imaging performance of the same sample. Note that we ensured to choose a highly transparent tissue (**Figure 6a inset)** for a fair comparison. As shown in **Figure 6 and Supplementary Video 5**, indeed LSTM allowed rapid uniform high-resolution imaging of the entire tissue, whereas LSM resulted in progressively poor image quality towards the middle of the sample, similar to the previous report(Stefaniuk et al., 2016). To complement these observations, we performed rapid high-resolution imaging of a large (3.32 cm x 1.93 cm x 1 mm) uniformly expanded (using Expansion Microscopy approach (Chen et al., 2015)) brain slice (**Figure 6c-d** and **Supplementary Video 6**). Expansion Microscopy (ExM) method is enabling unprecedented high super-resolution access to biological tissues, although also presenting unique challenges on imaging and data handling front. For example, imaging of this sample with the state-of-art Confocal or 2-photon microscopy will take several weeks of continuous imaging. We demonstrate unique suitability of LSTM for ExM samples by imaging (using 10x/0.6NA detection objective) the entire expanded tissue in ∼22 hours, yielding 723,200 images (2048x2048 pixels). The resulting dataset reveals the finest details of brain neuronal architecture (e.g. dendritic spines), while providing complete coverage. In addition, we also demonstrate imaging of a large piece of human brain tissue labelled with a uniform nuclear label DAPI. Human brain tissue, being dense, scatters the illumination light heavily and thus has been proven to challenging to be imaged by LSM approach. As shown in **Supplementary Figure 1**, LSTM indeed enabled uniform high-resolution imaging of large piece of human brain tissue.

**Figure 6.**
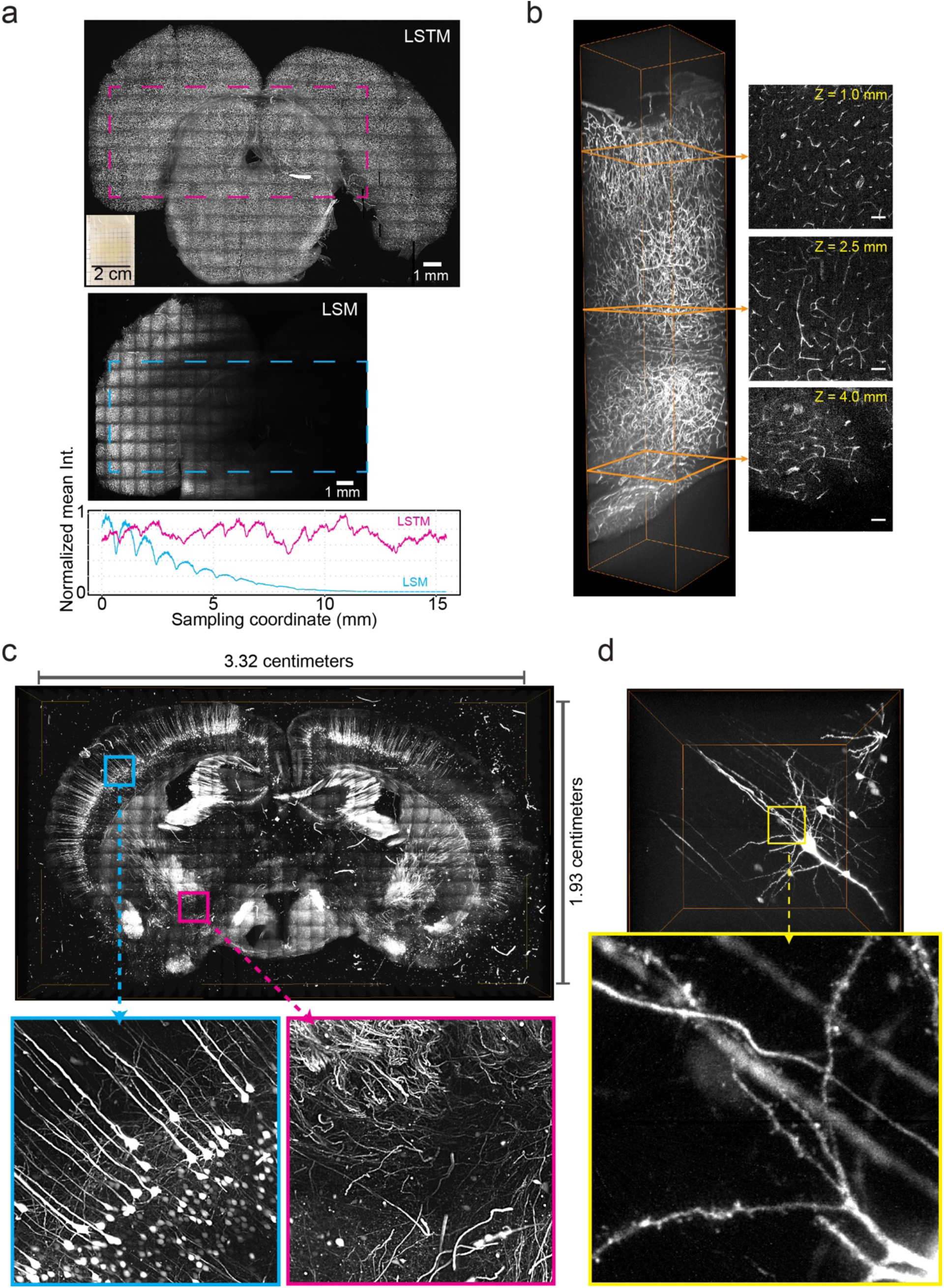
LSTM enables rapid uniform high-resolution imaging of very large samples. (a) For an unbiased comparison of the imaging performance of LSTM and LSM, we used a highly cleared large tissue sample (∼2 cm wide and ∼5 mm, inset in the top image) from rat brain. The sample was stained for visualizing the vasculature distribution. The images shown are maximum intensity z-projections, and were acquired using the same 10x/0.6NA objective. The bottom graph profiles the mean intensity (in the vertical direction) across the specified region of interest (ROI) marked with dotted rectangles. In LSM, the intensity signal is progressively degraded towards the interior of the sample, whereas LSTM allows uniform quality across the entire sample. The scale bars are 1 millimeter. (b) An example image stack from the LSTM dataset. Optical slices (50 microns thick) are shown at three different depths as identified (orange) in the stack. The bounding box of the stack is 1mm x 1mm x 5mm. The scale bars are 100 microns. A detailed volume rendering is shown in **Supplementary Video 5**. (c) Isotropically expanded (∼4-fold in all three dimensions) slice of *Thy1-YFP* transgenic mice was imaged using LSTM with 10x/0.6NA detection objective, resulting in 723,200 images (2048x2048). The imaging time was ∼22 hour. The shown volume rendering was performed with 8x8 fold down-sampled dataset. Zoomed-in images are shown as marked by colored rectangles and arrows. A detailed volume rendering is shown in **Supplementary Video 6**. (d) One image stack, from the dataset shown in c, is visualized. The bounding box size is 1.2mm x 1.2mm x 1mm. Note that, even with 10x/0.6NA objective, dendritic spines can be unambiguously identified. Imaging with 25x/1.0NA objective will result in further increased resolution, although at the cost of substantial increase in data size. See also **Supplementary Video 6** for detailed volume rendering.

Finally, we demonstrate the unique compatibility of LSTM in capturing the nervous system dynamics of a highly motile animal. Live samples often undergo substantial rearrangements in their body shape and cellular density, which significantly alter their local optical properties. Although LSM based imaging methods have been effective in capturing the cellular dynamics of developing embryos and neuronal activity of immobilized Zebrafish larvae, LSM remain susceptible to large changes in shape and density of motile sample, mainly because of the use of orthogonal illumination. This limitation has been partly addressed recently by utilizing a sophisticated array of hardware and software components that facilitate real-time adaptation of light-sheet parameters (Royer et al., 2016). LSTM, with its non-orthogonal illumination, may provide a simpler and highly effective solution. We tested this hypothesis by performing rapid volumetric calcium imaging of highly motile Hydra, which has been recently established as an effective model for exploring the role of neuronal network activities on apparent behaviors (Dupre and Yuste, 2017). We found that, indeed, LSTM enables aberration free calcium imaging of freely behaving Hydra which is undergoing drastic changes in its body shape and cellular density in the recording duration (**Figure 7a** and **Supplementary Video 7)**. In a way, the non-isomorphic body deformation of Hydra represents the worst-case scenario for tracking the activity of every neuron during behavior. We validated LSTM datasets by extracting and comparing neuronal traces with previous observations, finding excellent agreement (Dupre and Yuste, 2017). Note that, for this first demonstration, we used a relatively slow process of step-wise motion of the sample stage to acquire the image stacks. LSTM mechanism is straightforward to combine with piezo motor-based synchronous rapid scanning of the detection objective and also with extended detection depth-of-field(Tomer et al., 2015).

**Figure 7.**
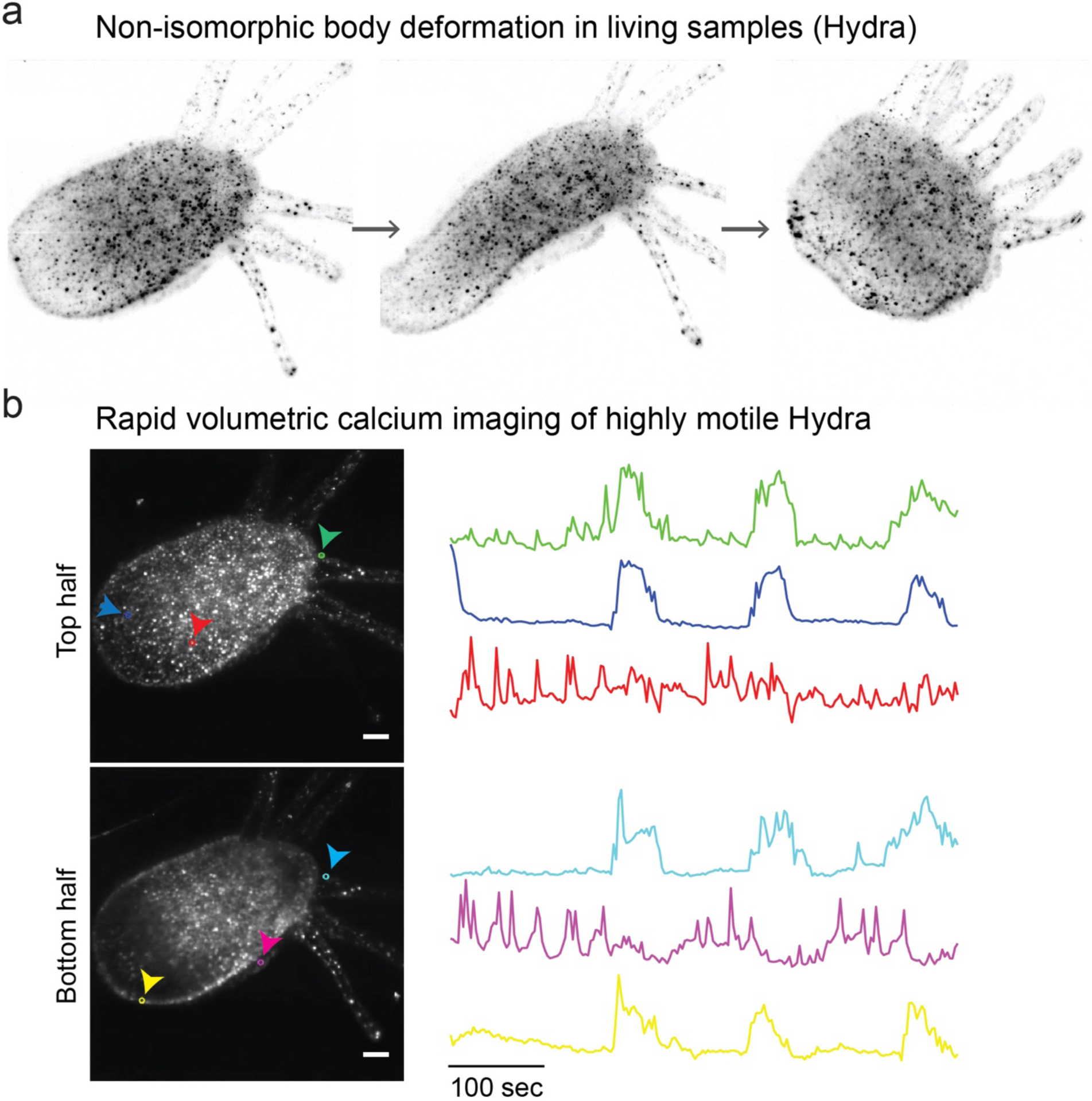
LSTM enables rapid volumetric imaging of highly motile animals. Live samples can undergo substantial non-isomorphic rearrangements in their body shape and cellular density, resulting in continuously changing local optical properties. LSM is particularly susceptible to misalignments and other aberrations because of the use of orthogonal light-sheet illumination. LSTM, by only using one-sided detection and illumination, is suitable for rapid volumetric live imaging of such difficult samples. (a) Hydra image is shown at different time points to highlight the non-isomorphic changes in freely moving animal. (b) LSTM was successfully used to perform long-term (>1 hour demonstrated, **Supplementary Video 7**) high-resolution live imaging of an adult Hydra expressing GCaMP6s(Dupre and Yuste, 2017). Each volume consisted of 17 z-planes. See **Supplementary Videos 7**-**8** for detailed visualization of the datasets. Manual tracking and analyses of calcium signaling was performed for the first ∼500 seconds of recording. Maximum intensity projections covering the two halves are shown. Representative neuronal traces are shown for cells marked in corresponding colors. As shown in **Supplementary Video 8**, four of the traces (plotted in green, blue, cyan and yellow) correlate with the rapid longitudinal contraction behavior of Hydra, and the other two traces are part of Rhythmic Potential circuits, in excellent agreement with the observations reported recently(Dupre and Yuste, 2017). Scale bars are 100 microns. See also **Supplementary Videos 7-8** for detailed volume rendering.

Thus, LSTM allows high-resolution quantitative imaging of large intact biological systems with no limitations on the lateral dimensions and high uniform quality. The sample thickness that can be imaged remains limited by the working distance of the detection objective and also by the level of tissue transparency and the penetration of labelling reagents.

## Discussion

Understanding the function of complex biological systems – especially highly complex mammalian brains - requires access to the intricate details of underlying structure and molecular composition, along with the functional dynamics. Over the last decades, a number of methods for clearing tissues to facilitate the interrogation of the structure and molecular architecture of large intact tissues have been developed. Methods such as CLARITY (Chung et al 2013, Tomer et al 2014) can render opaque tissues transparent while at the same time preserving structural and molecular content. Together with advances in high-speed imaging methods, these approaches provide new possibilities for understanding the functioning of tissues in health, and in diseased states such as malignant tumors and neurological damage. Methods that allow acquisition of high-resolution information on a practical time-frame such as LSM (Stelzer 2015, Migliori et al 2016) are effective for imaging these transparent tissues because of their inherent low photo-bleaching and the high imaging speeds.

The LSM approach of illuminating a sample with a thin sheet of light, and detecting the emitted signal with an orthogonally arranged detection arm provides two main advantages: minimal energy load and high imaging speed. We developed a highly-optimized implementation of LSM, called COLM(Tomer et al., 2015; Tomer et al., 2014) which allowed high-resolution imaging of entire intact mouse brains in a matter of hours. However, LSM as a general approach has been restricted in the lateral dimension of image volumes because the illumination light sheet enters from the side of the sample, and needs to penetrate the entire sample and the working distance of the illumination objective places a hard-physical limit of the imaging volume. In addition, physical tissue expansion approaches (Chen et al., 2015; Ku et al., 2016) for achieving high imaging resolution are producing even larger samples. To address these challenges, we developed a conceptually distinct imaging framework, called Light Sheet Theta Microscopy (LSTM). Similar to LSM, LSTM is based on planar illumination, but achieves this goal by using non-orthogonally arranged illumination objectives to produce light sheets that intersect the detection plane in a line profile, which is then synchronously (along with line-by-line detection of sCMOS camera) scanned along the detection plane. An immediate advantage of such a configuration is that it alleviates the restrictions on lateral sample dimensions, while providing uniform image quality for achieving true quantitative imaging. Note that LSTM imparts higher energy load (compared to LSM) for smaller samples, but similarly low photo-bleaching for larger samples. For all the imaging volumes reported in this study, no photo-bleaching is observed, further demonstrating the unique suitability of LSTM for imaging of samples of varied sizes and shapes. We found that a strategy of simultaneous 2-axes illumination (i.e. along and perpendicular to light sheet propagation) provided the best imaging performance. In comparison to LSM, LSTM allows imaging of larger samples such as a ∼2 cm wide and ∼5mm thick rat brain slice - with high uniform resolution across the entire sample. Moreover, LSTM is uniquely compatible with live imaging of highly motile samples, which undergo non-isomorphic changes in body shape and cellular densities.

LSTM is expected to allow uniform high-resolution imaging of large samples including thick slabs of cleared and labelled post-mortem healthy and diseased human brains as well as imaging of large animal intact brains, including rat and primate brains. Moreover, LSTM may facilitate *in situ* detection of thousands of transcripts in expanded tissue samples. Future work will include integration of super-resolution approaches (such as structured illumination) and simultaneous multi-view imaging.

## Methods

### LSTM implementation

The optical layout and physical implementation details are presented in **Figure 2, Supplementary Figure 2**. A complete parts list and their sources are listed in the **Supplementary Table 1**. In the LSTM implementation, two thin light sheets are generated by using two illumination optical arms each containing LASER source, cylindrical lens, vertical slit, iris, electrically tunable lens, galvo scanner, scan lens, tube lens, mirror, and the illumination objective (Olympus Macro 4x/0.28 NA). Since the illumination objectives were air objectives, we used a 3D printed cap (using Ultimaker 2+extended), containing a 1 inch diameter quartz coverslip, to seal the objective for oil immersion use. The emitted signal is detected by a detection arm which is composed of a detection objective, tube lens, and sCMOS camera (Hamamatsu Orca Flash 4.0 V3). The illumination arms were mounted at an approximately 60° angle relative to the detection arm on a vertically mounted breadboard fixed on an optic table with pillar posts. To facilitate the optical alignment of the system, all the three optical arms were mounted on two manual translation stages which were attached to the breadboard. We used a 3D printed (using Ultimaker 2+extended) open top sample chamber that was filled with an immersion oil of Refractive Index 1.454 (Cargille labs). The sample was mounted in a quartz cuvette (FireFly Scientific) which was then attached to the base of the sample chamber (**Supplementary Figure 2**). The 3D model of LSTM microscope was made with Autodesk Inventor 2017.

### LSTM geometric constraint calculations

The physical geometric constraints on the arrangement of illumination objectives were calculated by analyzing the two opposite bounds (**Figure 3a, Supplementary Figure 3**): the illumination objective not to touch the detection objective (**Figure 3a top**) and the illumination objective not going below the physical extent of the detection objective (**Figure 3a middle**). The range of allowable angular positions was calculated by taking the effective working distances and the objective diameters into account as shown in the schematics of **Supplementary Figure 3**, resulting in the following relationships:

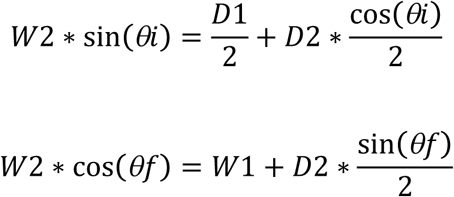

where W1 and W2 are the working distances of the detection and illumination objectives respectively, D1 and D2 are the diameters of the detection and illumination objectives respectively, and θi and θf are the boundary angular positions, as shown in **Figure 3a**, Supplementary Figure **3**. We used Macro 4x/0.28NA/29.5WD (Olympus) as the illumination objectives. These objectives are designed to be used in air, therefore we used Snell’s law to calculate the approximate effective working distance in a refractive index liquid of 1.454, as shown in **Supplementary Figure 3**, resulting in ∼44 mm, and the objective diameter (lowest part of the tapered ending) was measured to be ∼28 mm. For most of the experiments we used 10x/0.6NA/8mmWD objective (Olympus) in the detection arm with values of W1 = 8, and D1 = 40. For this combination, we found the allowable angular range to be 43.3° to 62.3° degrees. This calculated range was our initial guideline for identifying the maximum possible angular positioning during system alignment. We used ∼60° as the final angle separation. All calculations were performed in Matlab.

### Illumination depth calculations

We used geometric calculations (**Figure 3b**) to estimate the maximum illumination path lengths of LSTM as *t/* cos(*θ*), where t is the sample thickness to be imaged and θ is the angle between the illumination propagation direction and the detection axis. The maximum illumination path length in LSM would be the same as the sample width (w). We calculated the ratio of these illumination path lengths which was converted into a binary representation by thresholding at 1, and plotted as a heat map, shown in **Figure 3b**.

### LSTM effective light sheet thickness calculations

Due to the non-orthogonal incidence of the light sheet on the detection plane, the effective light sheet thickness can be approximated as the projection of the original thickness on to the detection direction, resulting in *b/*sin(θ), where b is the original light sheet thickness at the most focused position, and θ is the angle of incidence relative the detection axis. The relationship was plotted in the graph shown in **Figure 3c**.

### Imaging experiments

We used the passive CLARITY method (as described previously(Tomer et al., 2014)) for all the tissue clarification experiments. The hydrogel monomer (HM) solution recipe consisted of 1-4% (w*t/*vol) acrylamide, 0.05% (w*t/*vol) bisacrylamide, 4% paraformaldehyde, 1x Phosphate Buffered Saline, deionized water, and 0.25% thermal initiation VA-044(Wako Chemicals, NC0632395). All animal procedures were followed according to institutional guidelines.

For whole brain clearing, transcardiac perfusion was performed with 20 ml HM solution, followed by overnight incubation at 4°C. The rat brain was perfused with 4% paraformaldehyde (PFA), post-fixed for 16 hours and then frozen in isopentane for storage. The frozen brain was thawed at room temperature in PBS buffer, sliced and incubated in HM solution overnight at 4°C.

The human brain tissue was incubated in 4% PFA for ∼2 days, followed by incubation in HM solution up to days at 4°C. All the perfused tissues were de-gassed and transferred to 37°C for 3-4 hours for hydrogel polymerization. The tissues were cleared by incubating (with shaking) in clearing buffer (4% (w*t/*vol) SDS, 0.2 M boric acid, pH 8.5) at 37°C until clear (2-3 weeks). The rat brain and human brain slices were incubated in the HM solution overnight at 4°C followed by degassing to replace the oxygen with nitrogen, and incubation at 37°Cfor 3-4 hours for polymerization. After washing off any remaining HM solution, the tissues were incubated in buffered clearing solution (4% (w*t/*vol) SDS, 0.2 M boric acid, pH 8.5) at 37°C with shaking until the tissue became clear (1-4 weeks). Afterwards, the tissue was washed with 0.2 M boric acid buffer (pH 8.5) with 0.1% Triton X-100 for up to 24 hours. The cleared tissue was labelled with DAPI (1 μg/mL final concentration) and/or blood vessels marker tomato lectin (Vector Labs, FL-1171), by incubating in the labelling solution for 3-4 days. After washing with the buffered solution (0.2 M boric acid buffer, pH 7.5, 0.1% Triton X-100), the tissue was transferred into 85-87% glycerol solution in graded fashion (i.e. 25%, 50%, 65% and finally 87%) for final clearing and imaging. All the image volumes were acquired with a 2 or 5 microns step-size.

For isotropic tissue expansion, a *Thy1-YFP* mouse brain slice (250 um, perused and fixed with 4% paraformaldehyde and sliced with vibratome) was gelled and digested following the protein retention expansion microscopy (proExM) protocol(Tillberg et al., 2016). The sample was stored in 1x PBS before changing the buffer to 65% Glycerol (with 2.5 mg/mL DABCO) for the LSTM imaging.

For live imaging of Hydra, we followed the procedures described previously(Dupre and Yuste, 2017). Transgenic Hydra expressing GCaMP6s in neurons(Dupre and Yuste, 2017) were maintained in the dark at 18°C and were fed freshly hatched *Artemia nauplii* once a week, or more frequently when the colony needed to grow. Animals were mounted between two coverslips (VWR cat# 89015-724 and VWR cat#16004-094) using a 100um spacer (Grace Bio-Labs cat#654006) and were imaged with LSTM using 10x/0.6NA objective (Olympus).

### Image analyses

A TeraStitcher (Bria and Iannello, 2012) based pipeline(Tomer et al., 2014) was used for stitching of acquired image stack tiles of all the datasets. Maximum intensity projections, and other linear image contrast adjustments were performed using Fiji (Schindelin et al., 2012; Schneider et al., 2012) and MATLAB. All volume renderings were performed using Amira (FEI). All the fluorescent beads image analysis was performed using Fiji. To calculate the axial Full Width at Half Maximum (FWHM), x-z projections of beads image stacks were used. For individual beads a line intensity profile was calculated along the central position, followed by manual calculations of full width at half maximum intensity values. For Hydra data analyses, manual tracking of selected neurons was performed using TrackMate(Tinevez et al., 2017), and the generation of graphs and movies was performed using MATLAB.

### Illumination energy load calculations

The procedure is summarized in **Supplementary Figure 5**. To calculate the total illumination energy load in LSTM, we performed a simulation of the step-wise scanning of sample through the illuminating light-sheet. A horizontal plane across an entire sample can be imaged with approximately non-overlapping thin sheets of light, therefore, the total energy is calculated by step-wise scanning of the sample through the illumination volume. All voxels receiving illumination are incremented by 1. The final energy load is calculated as the total sum of accumulated illuminations in all voxels. The procedure was implemented of a range of parameters and two detection objectives (10x/0.6NA/8mmWD and 25/1.0NA/8mmWD). Each voxel in LSM imaging is illuminated w/f times, where w is the width of the sample, and f is the field-of-view size of the detection arm. Therefor the total energy load is ∼(w/f)*number of voxels. Note that the energy load in LSM as well as LSTM scales up by the same constant factor in LSTM and LSM, which cancels out in ratio,

### Data availability statement

All the datasets reported in this paper, ranging in tens of terabytes will be made available on request. Complete CAD model of LSM and other related resources will be made available with the manuscript and on a dedicated resources webpage. Complete parts list is included as a supplementary table.

### Materials & Correspondence

requests to Raju Tomer.

## AUTHOR CONTRIBUTIONS

R.T. conceived the project and designed the microscopes. B.M., M.S.D and R.T. built the microscopes. M.S.D., B.M. and R.T. performed the imaging experiments. M.C.A. assisted with all the experiments. C.D. and R.Y. prepared the Hydra samples for live imaging experiments. S.S., R.G. and E.S.B. generated expanded tissue. O.H. supported B.M. R.T., B.M. and M.S.D. analyzed the data. and wrote the paper with inputs from all authors. R.T. supervised all aspects of the work.

### ACKNOWLEDGEMENTS

We thank Weijian Yang for discussions and advice on the ETL use. We are grateful to Darcy Kelley and Oliver Hobert for general advice and reading of the manuscript draft. We would like to thank Serge Przedborski for providing rat brain tissue and Peter Canoll for the human brain tissue. This work is supported by Columbia University Arts & Sciences startup grant to R.T.. R.Y. and C.D. were supported by the NEI (DP1EY024503). The Hydra experimental work is based upon work supported by the Defense Advanced Research Projects Agency (DARPA) under Contract No. HR0011-17-C-0026.C-0026.

